## Supplemental Figures-Epigenetics Gene Families for "Epigenetics of skeletal muscle-associated genes in the *ASB, LRRC, TMEM*, and *OSBPL* gene families"

**Supplementary Figures**

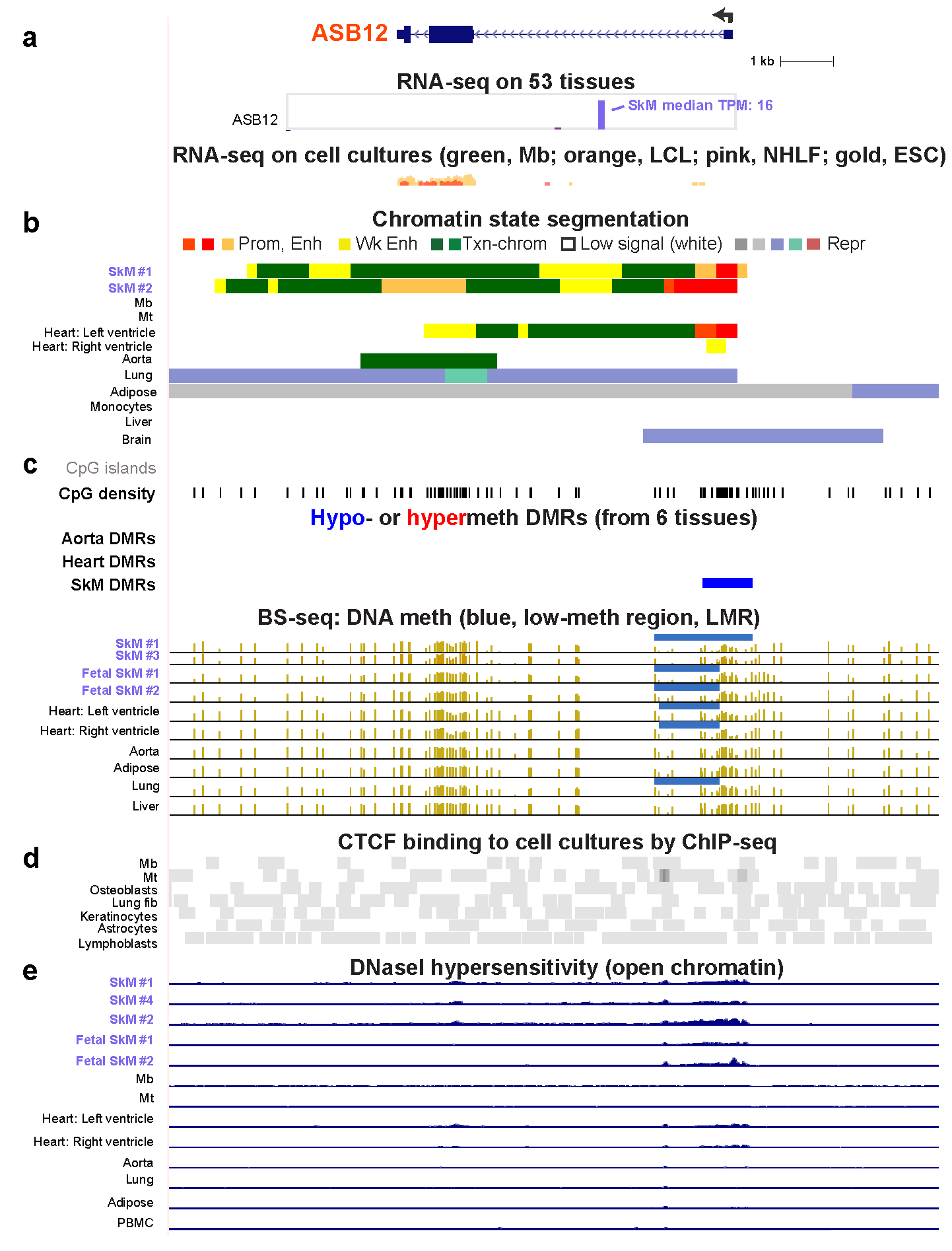

**Figure S1.** *ASB12* (ankyrin repeat and SOCS box containing 12), a little-studied SkM-specific gene, displays SkM-specific promoter-region hypomethylation like *ASB16, ASB4, ASB5*, and *ASB2*. **(a)** The RefSeq gene structure and RNA-seq analysis of expression of *ASB12* (chrX:63,439,714-63,454,446). (**b)** Chromatin state segmentation. (**c)** DNA methylation and CpG distribution. The CpG island track shows no signal indicating that an absence of CpG islands although the track for CpG dinucleotides shows a moderately high density of CpGs in the upstream promoter region. Panels (**a** – **c)** and (**e)** are notated as described in the main figures except that DMRs are shown for heart and aorta as well as for SkM with the six tissues being compared for determination of these DMRs as follows: SkM (psoas), heart (left ventricle, aorta, adipose, lung and monocytes. (**d)** CTCF binding from the UCSC Genome Browser. (**e)** DNaseI hypersensitivity (vertical viewing range, 0 – 10).

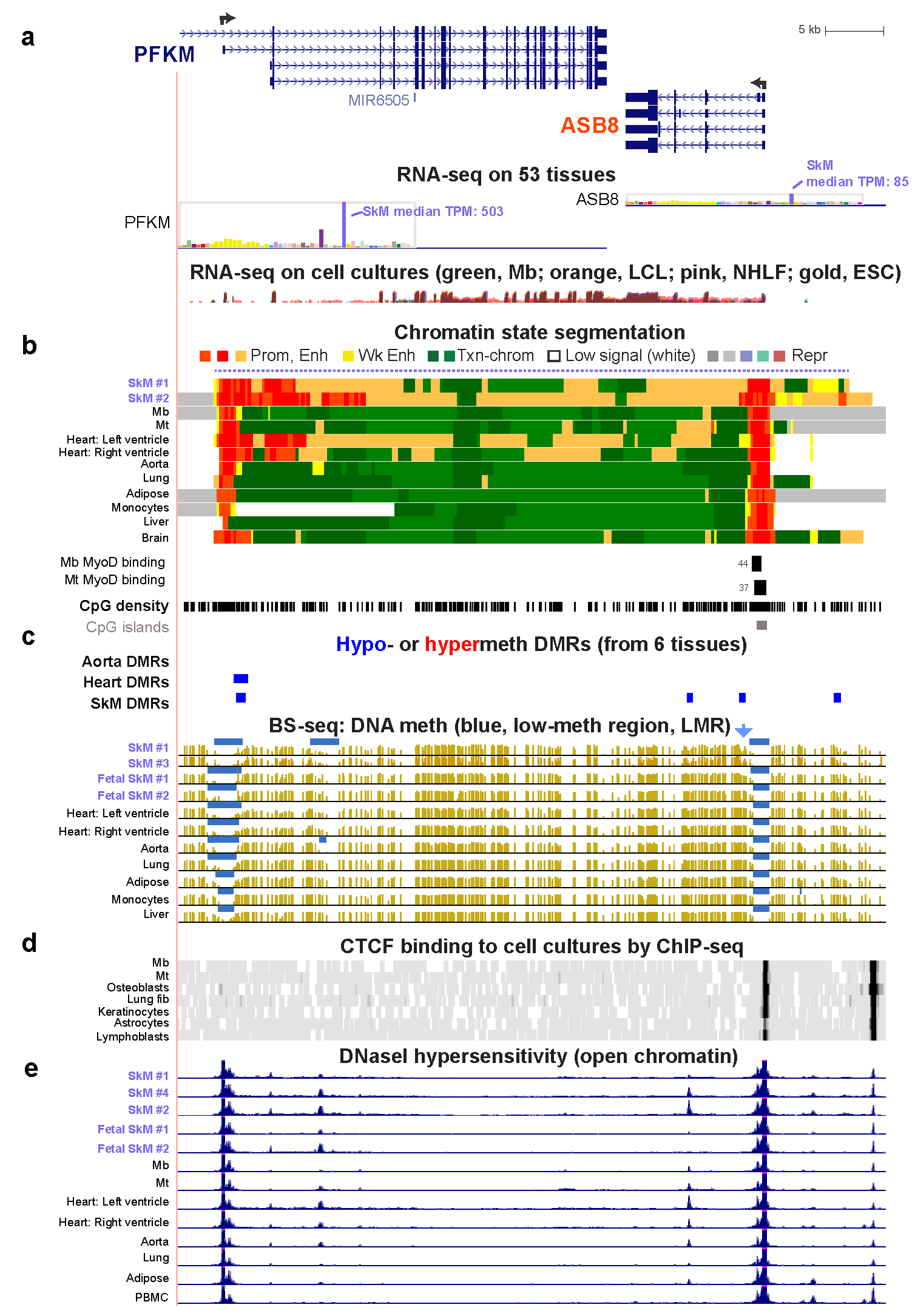

**Figure S2.** *ASB8* (Ankyrin repeat and SOCS Box Containing 8), and its adjacent gene *PFKM* (Phosphofructokinase, Muscle) are expressed at the highest levels in SkM and share a super-enhancer. **(a** – **e)** (chr12:48,509,874-48,559,915) are as described for Figure S1 except that the Mb-associated and Mt-associated mouse cell culture-deduced MyoD-binding sites and their signal in ChIP-seq are shown in panel (**b**). DNaseI hypersensitivity: vertical viewing range 0 – 30; dotted line in (**b**), SkM-specific super-enhancer.

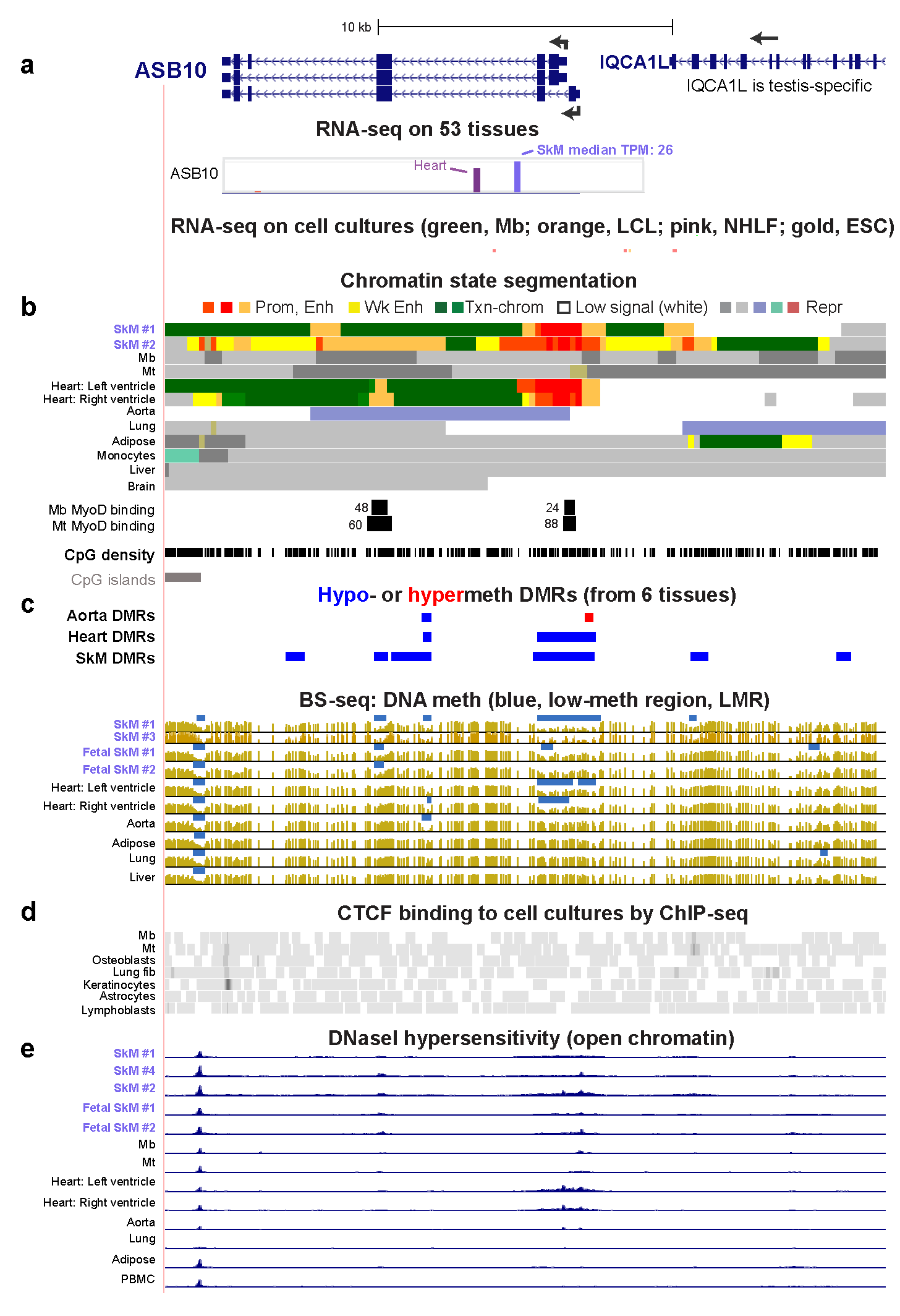

**Figure S3.** *ASB10* (Ankyrin repeat and SOCS Box Containing 10), which is preferentially expressed in SkM and heart, has an overlapping SkM/heart-specific enhancer-and-promoter chromatin containing SkM and heart hypomethylated DMRs**.** Panels **(a** – **e)** (chr7:150,861,978-150,895,455) are as described for Figure S2.

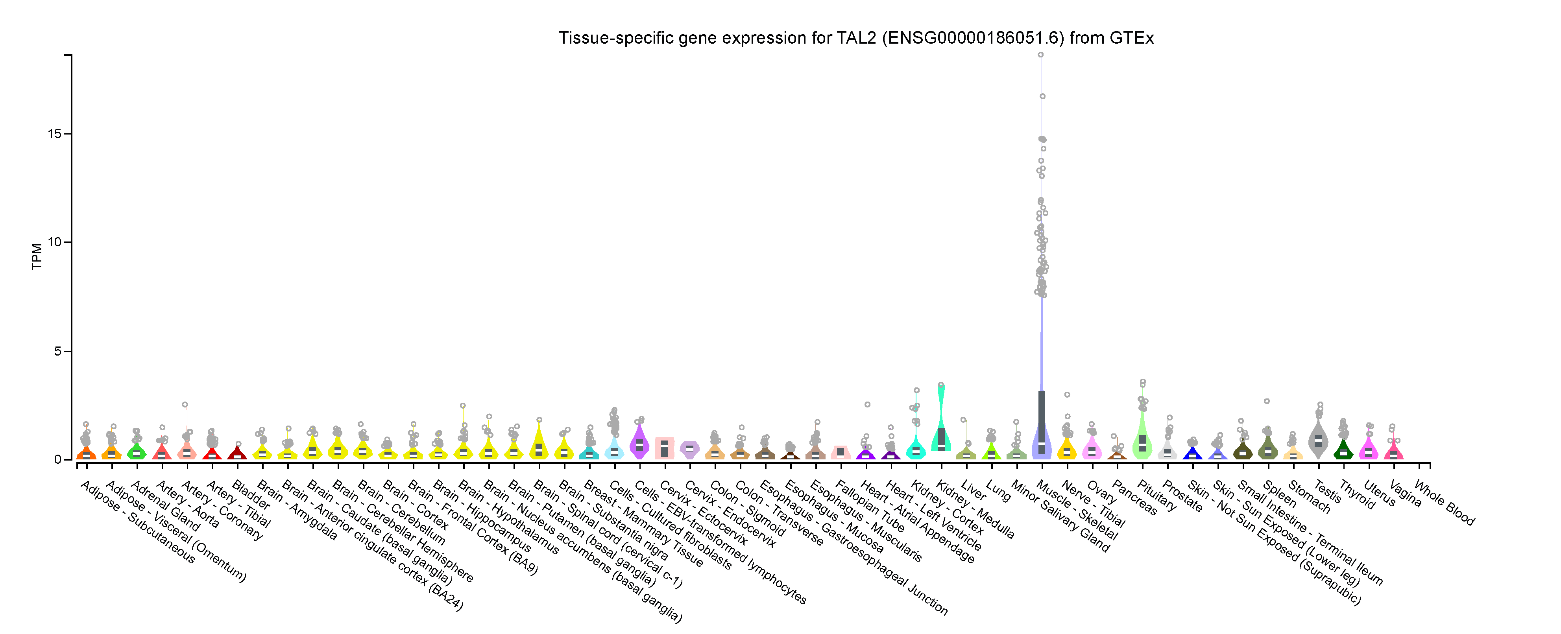

**Figure S4.** Detailed GTEx results for *TAL2* expression levels. Median levels of expression from hundreds of sample for each tissue type as determined by RNA-seq are shown.
